# DM: a simple solution to suppress air-water interface interactions in cryo-EM

**DOI:** 10.64898/2026.04.02.716008

**Authors:** Maria Rafiq, Jan-Hannes Schäfer, Hamidreza Rahmani, Shaochen You, Michael Bollong, Danielle Grotjahn, R. Luke Wiseman, Gabriel C. Lander

## Abstract

The air–water interface (AWI) remains the primary barrier to routine high-resolution cryo-EM structure determination, driving protein adsorption, structural denaturation, and restricted particle orientations during vitrification. Here, we describe a simple and broadly applicable strategy to mitigate these effects using the mild non-ionic detergent n-decyl-β-D-maltopyranoside (DM). Addition of DM at low millimolar concentrations immediately prior to vitrification consistently suppresses AWI-driven artifacts, resulting in improved angular sampling, reduced structural damage, and enhanced reconstruction quality across diverse macromolecular systems. Using this approach, we obtained a high-resolution reconstruction of the 65 kDa Nucleophosmin 1 pentamer, a target previously limited by severe preferred orientation issues. We further show that DM promotes isotropic particle distributions for high-resolution reconstruction of hemagglutinin, transthyretin, as well as suppressing denaturation of aldolase while stabilizing its C-terminus. Our results indicate that DM effectively passivates deleterious air-water interface interactions without compromising particle integrity. These results establish DM as an effective additive for improving the robustness of single-particle cryo-EM sample preparation.

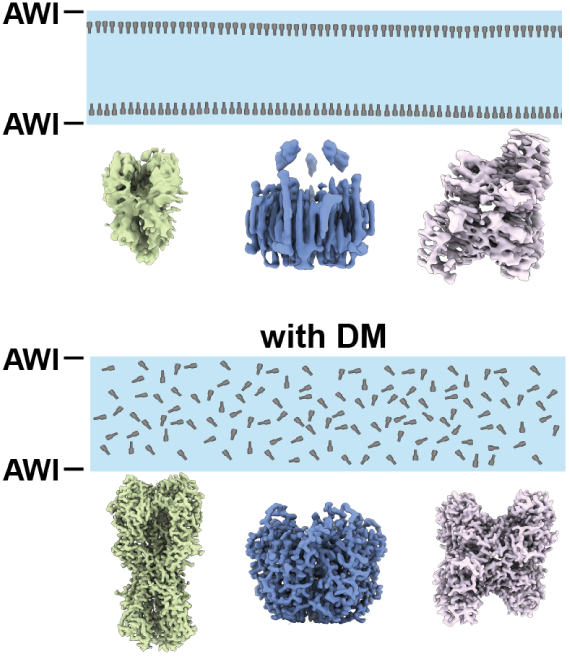

## Introduction

A persistent limitation in single particle cryo-electron microscopy (cryo-EM) is the behavior of macromolecules within the nanometer-scale aqueous films formed during sample vitrification. As samples are blotted, proteins diffuse rapidly and collide repeatedly with the air-water interface (AWI), a high-energy surface known to perturb structure and restrict accessible orientations. Cryo-electron tomography studies demonstrated that, for a majority of samples, 90-95% of particles localize within 10 nm of the AWI, rather than residing uniformly in bulk ice ^1^. The underlying physical basis for this phenomenon (surface tension, hydrophobic patches on the macromolecule, and partial unfolding at the AWI) are well-established and collectively position AWI exposure as a major determinant of single-particle reconstruction quality ^2–4^.

Proteins reach the AWI on the microsecond/millisecond timescale, far faster than the dwell times associated with conventional plunge-based vitrification ^5,6^. Once adsorbed, macromolecules often undergo partial unfolding, exposing internal hydrophobic residues that exacerbate AWI affinity and structural perturbations ^7^. Often, concurrent with these perturbations is a uniform adoption of particle orientation relative to the AWI – a “preferred orientation” – which restricts the angular sample of the targeted macromolecule and results in reconstructions with directional resolution anisotropy. Consequently, numerous strategies have been developed in an attempt to mitigate AWI adsorption ^5,8,9^, although none have emerged as the broadly applicable “silver bullet” for overcoming this prevalent issue.

Among the most widely used AWI interventions are detergents and surfactants, which compete for the AWI, shielding macromolecules from direct exposure to the hydrophobic surface. Non-ionic detergents such as DDM, GDN, OGNG and LMNG, as well as zwitterionic surfactants such as CHAPSO, have been reported to reduce AWI interactions by forming protective hydration layers ^10–12^. Amphipols similarly suppress AWI-induced membrane protein denaturation ^7,13,14^, and chemical additives such as PEG, trehalose, and glycerol alter film viscosity and interfacial tension, providing additional routes to reducing AWI interactions ^1,10,12,13,15–19^. More recently, LEA proteins have been hypothesized to form a protective barrier at the AWI, demonstrating the ability to shield fragile macromolecular complexes from structural distortions ^20^. However, despite their utility with specific samples, no single additive has been shown to reliably prevent preferred orientation across the diversity of macromolecular systems.

During our single particle cryo-EM studies of the Nucleophosmin 1 (NPM1) pentamer, we encountered severe orientation issues that were refractory to standard AWI mitigation strategies. In our subsequent search for surfactants that reduce AWI interaction, we identified n-Decyl-Beta-Maltoside (DM) as an unexpectedly effective additive. Although DM is widely used as a membrane-protein solubilizing detergent, its potential role as a deliberate interfacial modulator has not been systematically examined. We demonstrate that low-millimolar DM, added immediately prior to vitrification, substantially reduces particle adsorption to the AWI and broadens angular sampling and reduces associated particle damage across multiple structurally distinct complexes. These findings position DM as a practical and broadly accessible tool for managing AWI-driven artifacts in single-particle cryo-EM workflows.

## Results

### DM reduces preferred orientation of NPM1 pentamers to enable structure determination

To benchmark potential AWI-modulating additives, we targeted the NPM1 protein, which oligomerizes to form a 65 kDa pentamer and has been shown to adopt a severely preferred orientation in vitreous ice ^21^. In our attempts to overcome the preferred orientation of purified NPM1 particles, we screened detergents spanning multiple chemical classes at their critical micelle concentrations, including the non-ionic detergents dodecyl-maltoside (DDM) and lauryl maltose neopentyl glycol (LMNG), the cationic detergent Fos-choline 8, and the zwitterionic sterol detergent CHAPSO (see **Table S2** for details). In parallel, we evaluated NPM1 behavior on graphene-supported grids, poly-L-lysine-coated grids, and polyethylene glycol (PEG) cross-linked grids. Consistent with prior observations for small (*<*100 kDa) specimens, tilted data collection did not substantially alleviate preferred orientation, likely due to increased effective ice thickness and reduced signal-to-noise at high tilt angles ^6,9^. Across all conditions, particles were well dispersed and yielded high-quality 2D class averages, but the particle views remained dominated by a single projection (**Fig. 1A**). The resulting reconstructions exhibited strong directional resolution anisotropy, evidenced by low conical FSC area ratio (cFAR) values (0.07) indicating persistent interaction with the AWI and preferred orientation (**Fig. 1A**). We further performed 30-degree tilted data collection, but the analyses suffered from increased effective ice thickness and reduced signal-to-noise at high tilt angles ^6,9^, and a high-resolution structure was not obtained.

**Figure 1.**
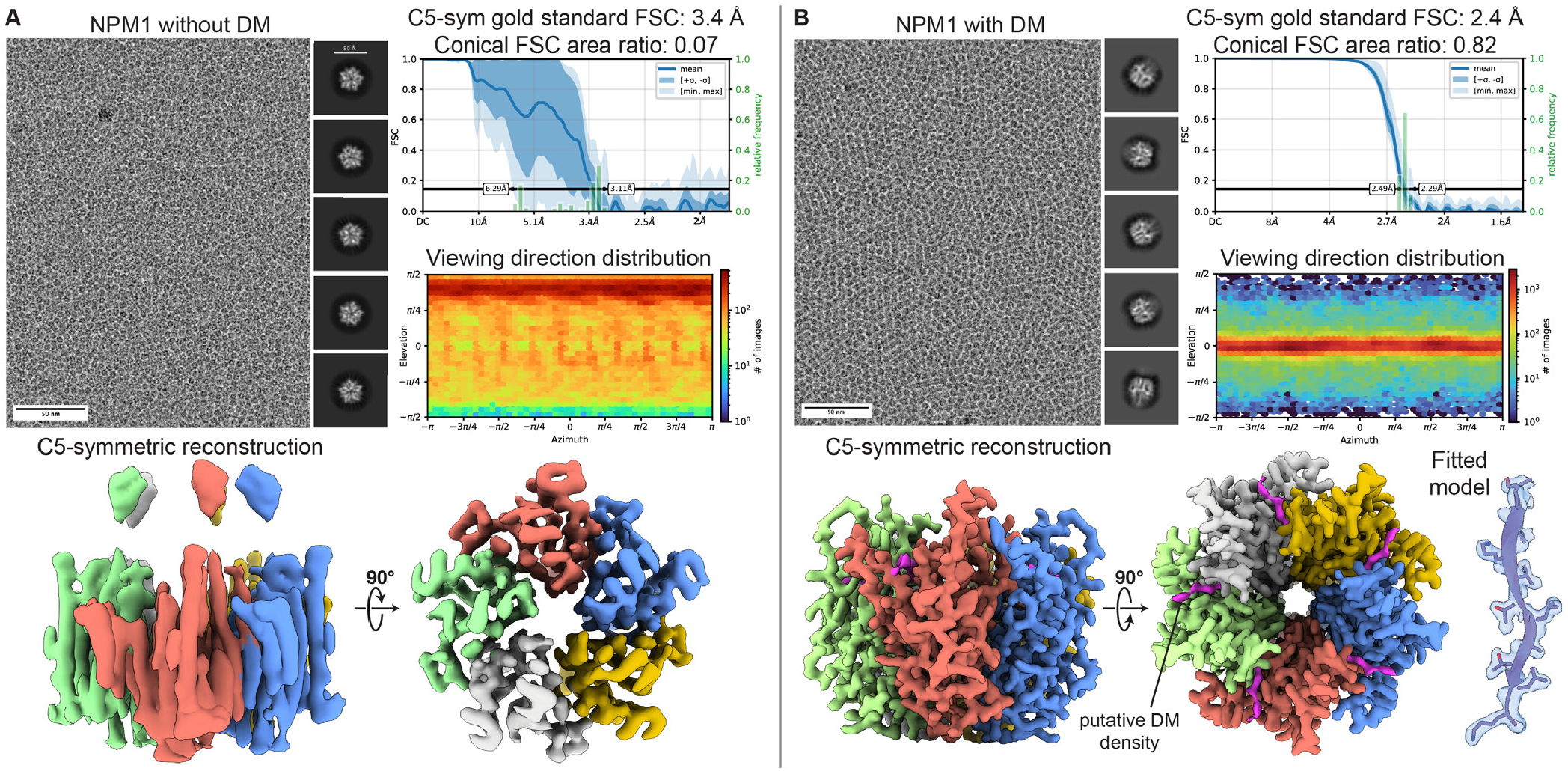
DM reorients Nucleophosmin 1 particles to overcome severe preferred orientation. (**A**) A representative micrograph and 2D averages of the NPM1 sample showing a single dominant view, consistent with strong preferred orientation. While the resolution of the reconstruction is 3.4 Å according to the FSC at 0.143, the conical FSC area ratio is low (0.07) and the directional distribution plots confirm preferential orientation assignment. Two orthogonal views of the resulting 3D reconstruction (lower panel) are shown with the surfaces colored by protomers, showing the hallmark “streaking” artifact associated with anisotropic directional resolution. (**B**) A representative micrograph and selected 2D averages of the NPM1 sample supplemented with DM shows that the majority of particles are now flipped to their side. The reported resolution of 2.4 Å is accompanied by a high conical FSC area ratio of 0.82. The direction distribution plots show that the dataset contains a full tomographic sampling of particle views around a single axis. Two orthogonal surface renderings of the resulting 3D reconstruction with directionally isotropic resolution (bottom left panel) are shown, colored by protomer, with putative DM density colored magenta. On the right, a close-up of residues 70-80 fit in their respective density demonstrate map quality.

We next tested if DM, a short-chain alkyl maltoside commonly used for membrane protein solubilization, could overcome the preferred orientation of NPM1 pentamers ^22–24^. DM was added to the NPM1 sample at its CMC (1.8 mM) immediately before vitrification, and the resulting micrographs showed monodisperse particle distribution without signs of aggregation or particle damage/dissociation. Notably, 2D analysis yielded class averages corresponding to views that were both distinct from the DM-free datasets and distinct from each other (**Fig. 1B**), indicating a substantial change in interaction with the AWI.

Quantitatively, the angular distribution broadened from a single dominant view in the asymmetric and C5-symmetric reconstructions (**Fig. 1A, Fig. S1**) to a continuous distribution spanning a single rotational axis (**Fig. 1B, Fig. S2**). Notably, DM did not fully randomize particle orientations but instead induced an 90º re-orientation within the ice, such that views orthogonal to the original preferred orientation became accessible. This redistribution enabled sampling across a previously missing angular axis and supported successful 3D reconstruction. Accordingly, the cFAR increased from 0.07 (in the absence of DM) to 0.82 with DM, reflecting attainment of the previously missing particle views. The resulting reconstruction was refined to a global resolution of 2.4 Å with a high degree of directional resolution isotropy, and uniform local resolution estimates across the core of the pentamer (**Fig. 1B**). Most importantly, side chain densities were qualitatively observed to exhibit details consistent with the reported resolution (**Fig. 1B**).

Intriguingly, the NPM1 reconstruction revealed additional elongated non-protein densities consistent with bound DM. These densities are located at inter-subunit interfaces on the surface of the pentamer, at the face of NPM1 that we previously observed to be oriented toward the air–water interface in the absence of DM (**Fig. 1B**). The residues comprising this putative DM-binding pocket are not strongly hydrophobic, suggesting that DM binding is not driven by a canonical hydrophobic interaction, but the interaction is nonetheless sufficient to alter local surface properties. We speculate that DM binding at this site contributes to the observed 90º reorientation of NPM1 particles within the ice, either by modulating surface hydrophilicity or by promoting interaction of an alternative surface of the particle with the AWI.

### DM broadly mitigates air-water interface interactions to enable structure

To determine whether DM’s effect was specific to NPM1 or more broadly applicable, we next examined the trimeric influenza hemagglutinin (HA), a protein assembly repeatedly documented to exhibit preferred orientation in vitreous ice ^9,25^. Addition of DM to the HA sample resulted in a broad coverage of views observed in 2D classes and in the angular distribution plots (**Fig. 2A**). With C3 symmetry imposed, HA trimer was resolved to a global resolution of 2.9 Å, with a cFAR of 0.85 indicative of an isotropic directional resolution profile (**Fig. S3**). As with NPM1, we also noted the presence of discrete elongated non-protein densities at the surface of HA map, which we ascribe to associated DM molecules (**Fig. 2A**, right).

**Figure 2.**
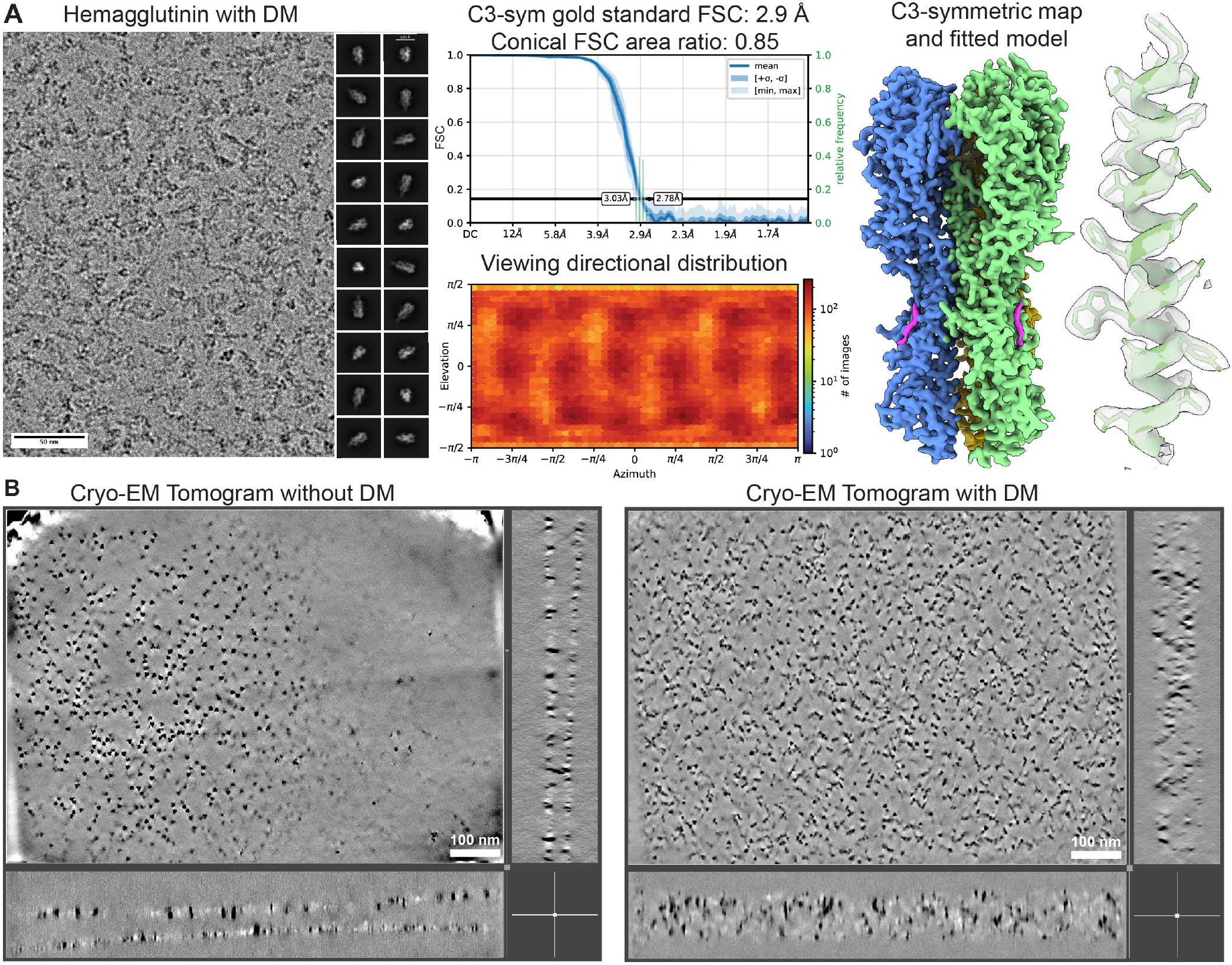
DM redistributes hemagglutinin and improves angular sampling. (**A**) Representative micrograph and 2D averages of the HA trimer prepared in the presence of DM. The structure was determined to have an overall resolution of 2.9 Å with a conical FSC analysis area ratio was 0.85, consistent with near-isotropic directional resolution. Directional distribution plots also support broad coverage across orientations. The 3D reconstruction is shown on the right as a surface representation colored by protomer. Discrete DM density is colored magenta are observed on the surface. On the far right a close-up of residues 75-100 in the EM density(transparent) demonstrates the quality of the reconstruction. (**B**) Cryo-electron tomography slices of HA grids prepared in the absence (left) and presence (right) of DM. For each condition, a central XY slice is shown, with corresponding orthogonal views (YZ slice, right; XZ slice, bottom) showing particle distribution in the vitrified ice.In the absence of DM, particles are predominantly located at the air-water interfaces, forming dense layers at the ice boundaries with minimal occupancy in the interior. In contrast, addition of DM results in a more uniform distribution of particles throughout the vitreous ice.

Due to the elongated shape of HA and its larger size relative to NPM1, we next aimed to directly assess HA particle localization within vitrified ice and its orientation within the ice using fiducial-free cryo-electron tomography. In the absence of DM, HA particles were confined to narrow layers at both AWI surfaces, leaving the interior of the vitreous ice sparsely populated (**Fig. 2B**, left). In contrast, tomograms of HA prepared with DM showed that the AWIs had effectively been reclaimed by the detergent (**Fig. 2B**, right), promoting a relatively uniform distribution of particles throughout the central section of the ice layer (**Movies S1, S2**). These data demonstrate that DM reduces interfacial trapping and promotes vitrification within the bulk ice, thereby restoring angular diversity in the imaged sample.

### DM stabilizes aldolase protomers by suppressing partial denaturation

Having shown DM’s ability to mitigate the preferred orientation of NPM1 and HA, we next tested whether DM could also reduce AWI-induced protein denaturation (**Fig. 3**). Rabbit-muscle aldolase was selected as a sensitive reporter system, as it has been shown to adhere to the AWI (Noble, Dandey, et al. 2018) causing partial denaturation of one of the subunits within the tetramer ^26^. In DM-free datasets, aldolase particles exhibited biased projection distributions and an asymmetric loss of density in one protomer in asymmetric reconstructions, presumably due to denaturing interactions of this subunit with the AWI (**Fig. 3C, Fig. S4**). Addition of DM increased the diversity of 2D class averages (**Fig. 3A**), broadened the angular distribution, improved directional FSC metrics, and restored density equally across all four of the protomers (**Fig. 3A-C**) in the D2 symmetric reconstruction as well as in the asymmetric C1 reconstruction (**Figs. S5, S6**).

**Figure 3.**
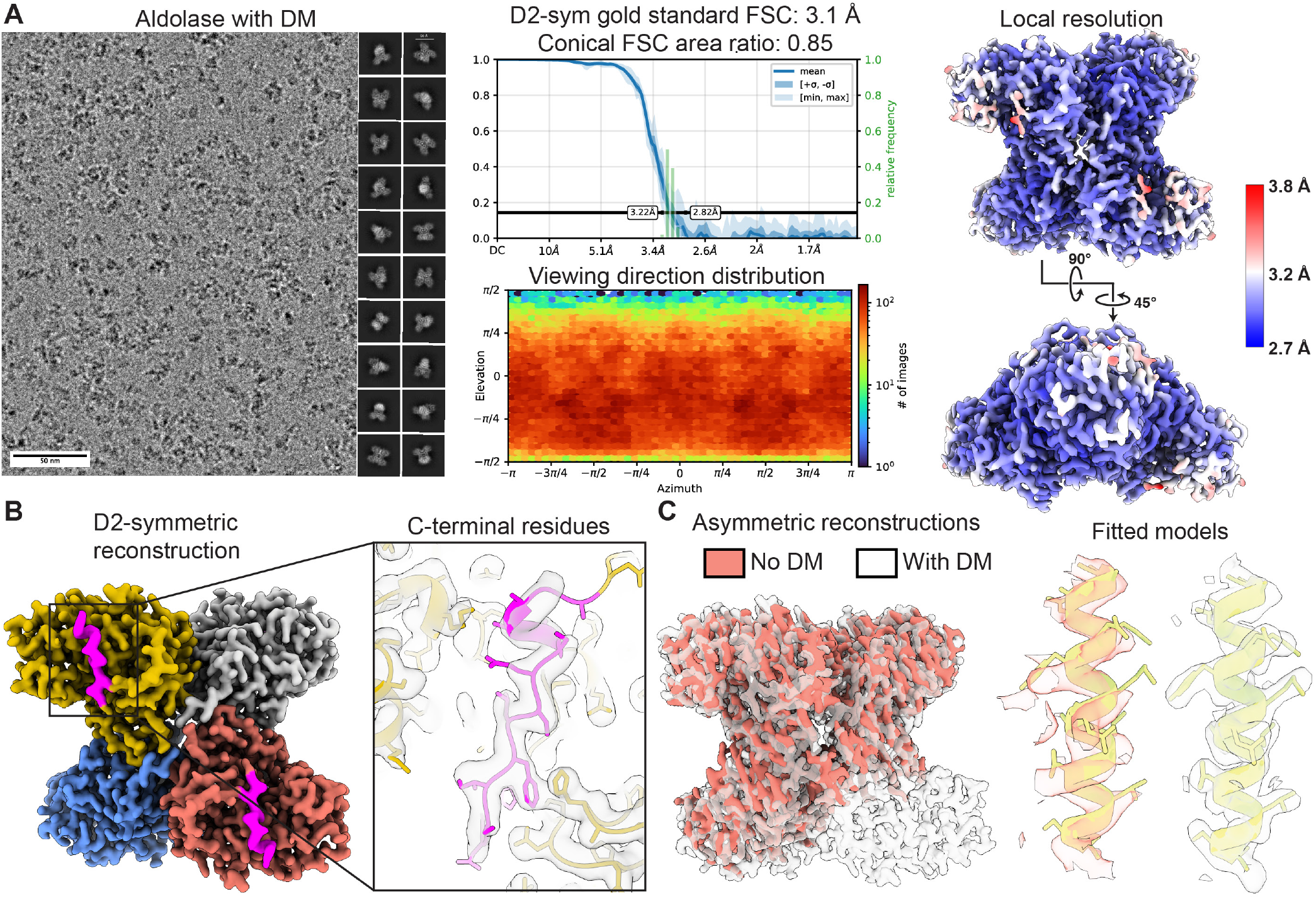
DM suppresses AWI-induced denaturation and preserves structural integrity of aldolase. (**A**) Representative cryo-EM micrograph and 2D class averages of aldolase prepared in the presence of DM, showing a broad range of particle views. To the right, conical FSC analysis (cFAR = 0.85) and viewing-direction distribution indicate near-isotropic angular sampling. On the far right, the aldolase map is shown colored by local resolution, demonstrates preservation of high-resolution features in peripheral domains. (**B**) Surface representation of the D2-symmetric reconstruction, colored by protomers. Density consistent with the C-terminal peptide is colored magenta. Inset shows a close-up of the density (transparent) with the atomic model of the aldolase C-terminus. (**C**) Comparison of asymmetric reconstructions of aldolase in the absence (red) and presence (transparent) of DM. In the absence of DM, one protomer exhibits a loss of ordered density, consistent with AWI-induced denaturation. In contrast, all four protomers are preserved in the DM-treated sample. On the far right a comparison of residues 321–339 from a preserved protomer in the detergent-free reconstruction and the corresponding region in the DM-treated dataset shows improved density with DM.

In the aldolase reconstruction, we observed additional density at the surface of each protomer that we initially attributed to bound DM. However, upon further inspection we found this density to be consistent with the C-terminal region of aldolase, which was structurally preserved in the presence of DM (**Fig. 3B**). To our knowledge, this region has not been resolved in prior cryo-EM reconstructions of aldolase. This observation indicates that DM efficiently suppresses AWI-associated structural damage, offering stabilization and capacity to visualize regions that might otherwise become disordered at the AWI.

### Applicability to smaller and larger assemblies: Transthyretin and ClpP protease

To further assess the ability of DM to mitigate air–water interface effects for smaller proteins, we examined the soluble 55 kDa homotetramer transthyretin, which exhibits pronounced preferred orientation that was previously resolved using graphene-supported grids ^27^. Consistent with our observations for NPM1, aldolase, and HA, addition of DM to the TTR sample prior to vitrification broadened angular distribution, yielding a substantially more diverse set of 2D class averages (**Fig. 4A**) and enabling a symmetric (D2) reconstruction with improved directional resolution with an estimated global resolution of ∼ 3.2 Å (**Fig. 4B, Fig. S7**). An asymmetric reconstruction of the homotetramer confirmed that none of the subunits were preferentially denatured (**Fig. S8**).

**Figure 4.**
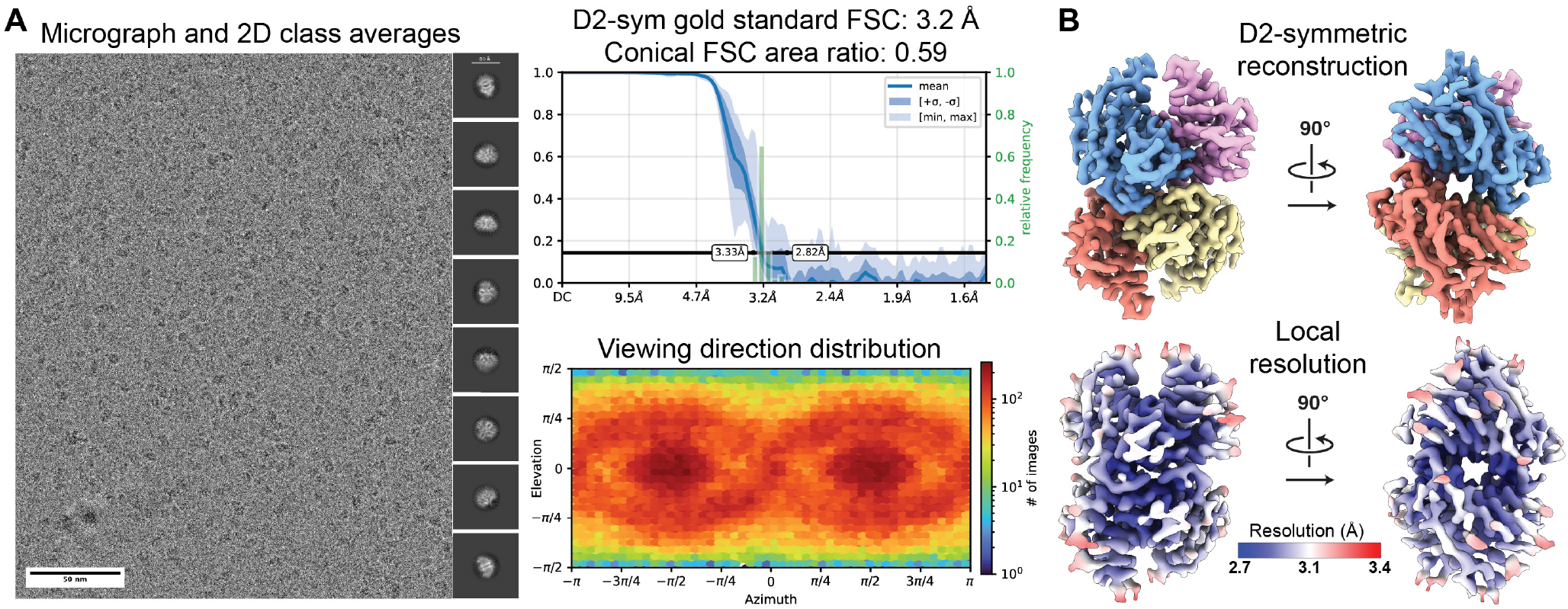
DM improves angular sampling for transthyretin. (**A**) Representative micrograph and 2D averages of transthyretin in the presence of DM. The global resolution is estimated to be 3.2 Å with a moderate level of directional resolution isotropy, as shown by the conical FSC analysis (cFAR = 0.59) and broad distribution of viewing directions. (**B**) Two orthogonal views of the final D2-symmetric TTR reconstruction are shown, colored by protomers (top), and estimated local resolution (below).

It should be noted that, despite improving the overall angular distribution of the particles in vitreous ice, the DM-containing sample retained some residual orientation bias, as evident in the directional sampling plot (**Fig. 4A, Fig. S7**). Notably, unlike the other systems examined here, we did not observe density attributable to bound DM in the TTR reconstruction, suggesting that its effect in this case is mediated primarily through weaker interfacial passivation rather than direct protein surface modulation.

These results indicate that while DM substantially improves sampling for a given macromolecular system, it may not fully eliminate preferred orientation in all cases, underscoring that no single additive universally resolves AWI-driven artifacts. Instead, these findings highlight the potential value of combining DM with complementary strategies, including multi-detergent approaches, as recently proposed ^28^, for particularly challenging targets.

During the preparation of this manuscript, we employed the new-found utility of DM to overcome the preferred orientation of the human ClpP protease complex ^29^. As observed for smaller soluble proteins, DM broadened accessible views and improved effective resolution along previously under-sampled axes without perturbing stoichiometry or domain organization. Consistent with our observations for other samples presented here, EM density attributable to DM was located on the ClpP map, at protein surfaces that would otherwise be interacting with the AWI in the absence of DM.

### Comparative conclusions from the detergent screen

Across multiple protein classes – from small symmetric tetramers, pentameric assemblies, and viral glycoproteins, to larger protease complexes - DM consistently outperformed other detergents tested under identical conditions. Whereas DDM, LMNG, Fos-choline 8, and CHAPSO failed to block the stubborn AWI-interface interactions of NPM1, DM restored isotropic distributions and enabled high-resolution reconstructions. The combined single-particle and tomographic data support a model in which DM modifies the interfacial free energy landscape at the AWI, reducing adsorption and suppressing both orientation trapping and partial denaturation. Importantly, its effectiveness extends across structurally and biochemically distinct systems. Taken together, this comparative screen identifies DM as a broadly effective and practical vitrification additive that can be readily implemented to mitigate AWI-driven artifacts in routine single-particle cryo-EM workflows.

## Discussion

Despite major advances in instrumentation and image processing, single-particle cryo-EM remains constrained by the behavior of macromolecules in thin aqueous films. During the interval between blotting and vitrification, macromolecules rapidly diffuse to the AWI, where adsorption can promote partial unfolding and trap particles in a restricted set of orientations ^1^. Because protein adsorption reduces the total interfacial free energy, the AWI provides a strong thermodynamic sink. Thus, reagents that lower surface tension or compete for the interface weaken the thermodynamic driving force for protein adsorption ^13,30,31^. This framework provides a unifying explanation for why certain surfactants or engineered supports reduce preferred orientation. Increasingly, specimen-preparation studies converge on a clear conclusion: for many targets, control of the AWI sets the upper limit on achievable resolution.

Chemical additives are among the simplest experimental routes for mitigating AWI-driven preferred orientation. Carefully chosen non-ionic detergents and polymeric surfactants tend to reduce adsorption and broaden orientation distributions for some targets ^10,32^ but can introduce additional heterogeneity and are not universally compatible ^14,33,34^. We thus face a sample screening challenge when we encounter a sample that exhibits strong interactions with the AWI, as no single reagent offers a broadly reliable solution that is effective across diverse samples.

Within this landscape, DM occupies an interesting niche. DM is a widely used non-ionic maltoside with favorable protein stabilizing properties and a relatively high CMC (∼1.8 mM) ^11^. Although DM supports high-resolution cryoEM studies of membrane proteins through its use in solubilization, purification, and reconstitution, DM is never systematically titrated around its CMC, nor has it been systematically investigated for its potential as an AWI-blocking additive. Our results establish DM as an effective and practical reagent for reducing preferred orientation and AWI-associated denaturation.

For all proteins tested (NPM1, aldolase, HA, TTR, and ClpP), adding DM immediately before vitrification resulted in markedly improved orientation distributions. DM broadened angular sampling, eliminated dominant views, and yielded more isotropic reconstructions (**Figs. 1-4**). Cryo-ET confirmed that DM displaces particles from both AWI surfaces, redistributing them into the interior of the ice layer (**Fig. 2B**). These improvements occurred without aggregation, oligomeric disruption, or detectable structural instability, consistent with DM’s profile as a mild detergent.

Interfacial studies of C10 maltosides suggest they form expanded monolayers near the CMC, reducing surface tension and lowering the energetic benefit of protein adsorption ^35,36^. This model is consistent with our cryo-ET observations. In addition, our observation of discrete non-protein densities consistent with bound DM in two of our test proteins suggest low-affinity or transient interactions that may partially mask hydrophobic regions and further disfavor AWI adsorption. Importantly, these interactions did not visibly perturb the protein structure.

In light of these collective observations, we favor a simple model for DM functioning at AWI (i) rapidly assembles into a dynamic monolayer at the AWI that competes with proteins for the interface, and (ii) transiently coats exposed hydrophobic patches on the protein, reducing direct protein interface contacts. (iii) mitigates protein denaturation by limiting AWI exposure through detergent adsorption at these interfaces. This role is consistent with broader work suggesting that effective AWI mitigation usually requires both interfacial engineering and subtle tuning of particle surface properties ^10,31,37,38^. DM, like other small-molecule additives, may interact with promiscuous active sites and potentially induce structural changes. Although no such effects were observed in our study, we recommend titrating DM to the lowest effective concentration to minimize potential artifacts. In our case, 0.05% (v/v) was sufficient to randomize orientations. Structural analyses performed in the presence of DM should be supported by activity assays under the same conditions to confirm preserved function.

Although no single approach universally solves AWI-driven preferred orientation, DM fills a useful niche. It is inexpensive, non-ionic, and already familiar to many laboratories. Screening DM near its CMC provides a low-effort strategy for samples that tolerate maltosides and exhibit severe preferred orientation, offering a straightforward alternative before more specialized methods are attempted. Our results demonstrate that low-millimolar DM reliably suppresses preferred orientation and AWI-induced denaturation across diverse macromolecules, establishing it as a practical and broadly applicable additive for improving cryo-EM specimen preparation.

## Material & Methods

### Protein expression and purification

#### NPM1

NPM1 was received as a gift from the Bollong Lab at Scripps Research. The protein was purified as described previously ^21,39^. Briefly, NPM1 9-122 construct (Addgene #23142) was expressed in *E. coli* BL21(DE3) cells (New England Bio-labs) in LB culture medium. After growth to OD_600_ around 0.6, protein expression was induced by 1 mM isopropyl b-D-thiogalactopyranoside (IPTG) and cells were incubated at 30°C for 18 h. The cells were lysed by sonication in lysis buffer (20 mM Tris-HCl pH 8.0, 500 mM NaCl, 5% v/v glycerol). The lysate was centrifuged at 18,000 X *g* for 30 min to remove insoluble material. Recombinant protein was purified first using nickel nitrilotriacetic acid agarose affinity chromatography (Ni-NTA, Qiagen), with 25 mM imidazole wash and 250 mM imidazole elution, then gel filtration on a Superdex 75 Increase 10/300 GL column (Cytiva) using FPLC buffer (20 mM Tris-HCl pH 7.5, 100 mM NaCl). Fractions corresponding to NPM1 were pooled and concentrated to 10 mg/ml for cryo-EM sample preparation.

#### HA-trimer

CA09 E47KHA trimer was received as a gift from the Ward Lab at Scripps Research. Protein expression and purification was performed as previously described ^40^. Briefly, HA proteins were transiently expressed in HEK 293F cells (Thermo Fisher) seeded at 1.0 x 10^6^ cells/mL and transfected at a 1:3 DNA to PEI-Max ratio. Cells were maintained in FreeStyle™ 293 expression medium (Life Technologies) at 37°C, 8% CO_2_, with shaking at 125 rpm. Six days post transfection, cells were harvested by centrifugation and HA was purified from the supernatant using a His Trap affinity column (Cytiva). To remove the His tag, HA protein was incubated with Tobacco Etch Virus (TEV) protease at a 1:20 molar ratio of TEV to HA for 16 hours at room temperature. Cleaved protein was further purified by size exclusion chromatography on a Superdex 200 Increase 10/300 column (GE Healthcare). Fractions corresponding to trimeric HA were pooled, concentrated up to 8 mg/ml and buffer exchanged into TBS using 50 kDa Amicon centrifugal concentrators.

#### TTR

TTR tetramer was received as a gift from the Kelly Lab at Scripps Research. The protein was expressed and purified as described in the previous work ^27^. Briefly, wt TTR was expressed in BL21 (DE3) *E. coli* transformed with the pMMHa vector encoding the wt *TTR* gene and selected with 100 µg/ml ampicillin. Cultures were grown to OD_600_ ∼ 0.5 at 37 °C and induced with 1 mM IPTG, followed by overnight expression at 30 °C. Cell pellets were resuspended in Tris-buffered saline supplemented with EDTA-free protease inhibitors and lysed by probe sonication (3 cycles, 3 min on/3 min off at 4°C). Clarified lysates were subjected to sequential ammonium sulfate precipitation at 50% and 90% (w/v), the pellets were collected and dialyzed against 25 mM Tris pH 8.0. The protein was purified by anion exchange chromatography (SourceQ15) using a NaCl gradient and further purified by size-exclusion chromatography (Superdex 75) in 10 mM sodium phosphate pH 7.6, 100 mM KCl. TTR was pooled and concentrated to 10 mg/ml for cryo-EM sample preparation.

#### Aldolase

Lyophilized rabbit muscle aldolase (Sigma-Aldrich) was solubilized in 20 mM HEPES pH 7.5, 50 mM NaCl to a final concentration of 8 mg/mL.

### Sample preparation for cryo-EM

NPM1, HA, and aldolase samples were applied to 300 mesh R1.2/1.3 or R0.6/1.0 UltrAuFoil Holey Gold Films (Quantifoil) that had been glow discharged for 25 s at 15 mA using a Pelco Easiglow 91000 (Ted Pella, Inc.) in ambient vacuum. All subsequently described vitrification steps were performed at 4 °C and *>*95% humidity using the same procedure. The concentrated samples of NPM1 (10 mg/ml) and aldolase (8 mg/ml) were supplemented with DM to a final concentration of 1.8 mM (0.08% v/v) immediately prior to grid preparation. 3.5 µL of sample was applied on grid, and excess sample was blotted away with Whatman No. 1 filter paper until the liquid spot on filter paper stopped expanding (approx. 5 s). The grids were manually plunge frozen in liquid ethane cooled by liquid nitrogen. For HA (8 mg/ml), DM was added at a final concentration of 0.06% (v/v) and vitrified in the similar way as described for NPM1 and aldolase. For TTR, the sample was concentrated up to 10 mg/ml, and 0.06% (v/v) DM was added immediately prior to vitrification. 3.5 µL of sample was applied to the grid, blotted for 4-6 s.

### Cryo-EM data acquisition

All data acquisition was performed on a ThermoFisher Talos Arctica transmission electron microscope operating at 200 keV and equipped with a Falcon 4i direct electron detector at a nominal magnification of 190,000, corresponding to a nominal pixel size of 0.74 Å and a calibrated pixel size of 0.731 Å . Movies were collected in counting mode, with a total exposure of 50 e^-^/Å ^2^, saved in the electron-event representation (EER) format. All datasets were collected using EPU v.3.9 with a defocus range set from -0.8 to -2.0 µm. Data was acquired at an 8 µm image shift using EPU fast exposure navigation. Movies were fractionated into 40 frames for processing, and all image analysis was performed in cryoSPARC (v.4.6-7) ^41^. Detailed processing workflows and dataset specific parameters are provided in the supplementary material and **Table S3**.

### Cryo-EM data processing

#### NPM1

The image processing pipeline for the detergent-free NPM1 structure is shown in **Fig. S1**. A total of 4,202 movies were collected and processed in cryoSPARC. Movies were motion corrected and dose-weighted using patch-based motion correction, and CTF estimated with cryoSPARC Live. A total of 3,129 micrographs with CTF fit values of 5 Å resolution or better were selected for further processing. The cryoSPARC blob picker was used to pick particles with a diameter of 80 Å, yielding approximately 3 million particles. Particles were extracted using a 216-pixel box size and binned to 100 pixels. 2D classes were generated using a circular mask of 80 Å, number of online iterations: 40, batch size per class of 200. 2D classes with 1.1M particles displaying high resolution secondary features of pentamer were selected. *Ab-initio* reconstructions were calculated using 2 classes and a resolution range of 8-35 Å . The resulting initial volumes were then used for heterogeneous refinement to classify particles into four 3D volumes with initial resolution 12 Å and spherical mask diameter of 90 Å . The 259,951 particles contributing to the best-resolved 3D class were re-extracted with the full box size (216 pixels) and subjected to a final round of non-uniform refinement using an initial low-pass filter of 8 Å, yielding a final asymmetric reconstruction with a global resolution of 8 Å, based on the FSC 0.143 cutoff criterion. Imposing C5 symmetry resulted in a reconstruction with a reported global resolution of 3.4 Å . However, the conical FSC plots for these reconstructions reveal severe directional resolution anisotropy.

The image processing pipeline for the NPM1 structure supplemented with DM is shown in **Fig. 2**. A total of 2,639 micrographs with CTF fit estimates of 5 Å resolution or better were selected after preprocessing. The cryoSPARC blob picker was used to pick particles with a diameter of 80 Å, yielding approximately 4 million particles. Particles were extracted using a 216-pixel box size, and binned to 100 pixels for preliminary analyses, beginning with classification into 50 2D classes. Classes were generated using circular mask 76 Å, number of online iterations: 40, batch size per class:200, and two rounds of 2D classes were performed to remove 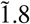 M junk particles. 2D classes with 1,204,896 particles displaying high resolution secondary features of pentamer were selected to compute two reference free 3D *ab-initio* models. The resulting initial volumes were then used for heterogeneous refinement to classify particles into three 3D volumes with C5 symmetry imposed and initial resolution 10 Å and spherical mask diameter of 90 Å . The particles from the best class were re-extracted with the full box size (216 pixels) and subjected to a final round of non-uniform refinement with C5 symmetry using an initial low-pass filter of 5 Å . A mask focused on the pentamer was generated in ChimeraX ^42^ by using the “molmap” command, imported into cryoSPARC and converted into a mask using 0.0575 threshold and 10-pixel soft padding. Local refinement was done with a generated mask on pentamer, with rotation search extent 1 degree and shift search extent 1 Å, an initial low-pass resolution filter of 4 Å, yielding a final reconstruction with a global resolution of 2.4 Å based on the FSC 0.143 cut off criterion.

#### Hemaggluttinin

The image processing pipeline for the HA analysis is shown in Fig. S3. A total of 3,542 movies were collected, dose-weighting, motion correction, and CTF estimation were performed in cryoSPARC Live. Particles were picked using blob picking with a particle diameter of 200 Å, yielding approximately 728,939 particles. 2D classification into 50 classes with default parameters were used to remove junk, and well resolved classes containing 614,028 particles were carried forward into two *ab-initio* reconstructions. In parallel, heterogeneous refinement with force hard classification true and batch size of 2000 per class into four classes was performed on the full set of cleaned 614,028 particles. The class with the best resolved structural features, comprising 325,210 particles, was subjected to non-uniform refinement without symmetry, with a 12 Å lowpass-filtered input and “minimize over perparticle scales” enabled, yielding an HA reconstruction at 6 Å resolution. Particles were then re-extracted at the full box size of 400 pixels and subjected to non-uniform refinement, with a 12 Å lowpass-filtered input and “minimize over per-particle scales” enabled followed by global and local CTF refinement for three iterations and a final round of non-uniform refinement with C1 symmetry, producing a map with a reported gold-standard FSC global resolution of 3.1 Å . After applying C3 symmetry the resolution improved to 2.9 Å, based on the FSC 0.143 cutoff criterion.

#### Aldolase

The image processing pipeline for the detergent-free aldolase dataset is shown in **Fig. S4**. We reprocessed a dataset previously acquired by our group for another study ^43^. Briefly, 1.4 mg/ml aldolase was used to prepare grids, particles were picked with the blob picker in cryoSPARC Live using a particle diameter of 100-110 Å . 820,058 particles were extracted with a box size of 200 pixels, Fourier cropped to 100 pixels, and 2D classification was performed requesting 50 classes. Subset particle selection was performed to yield 389,144 particles. *Ab initio* reconstruction was performed requesting two classes using C1 symmetry and an initial resolution set to 8 Å . These initial volumes were used as input for a heterogeneous refinement job using all particles, where 415,330 particles classified into a well-resolved class structurally consistent with the aldolase tetramer. These particles and density were next further refined with non-uniform refinement using C1 symmetry, with the initial low-pass resolution set to 8 Å and per-particle defocus and per-group CTF parameter optimization turned on. These particles were re-extracted with a box size of 280 pixels and further refined using non-uniform refinement with C1 symmetry, an initial low-pass resolution set to 8 Å, and per-particle defocus and per-group CTF parameter optimization turned on, yielding a reconstruction with a reported resolution of 3.0 Å based on an FSC cutoff of 0.143.

The image processing pipeline for the aldolase structure in the presence of DM is shown in **Fig. S5**. A total of 750 micrographs were collected and dose weighting, motion correction, and CTF estimation were performed in cryoSPARC Live. The cryoSPARC Blob picker was used to pick particles with a diameter of 150 Å, yielding approximately 605,114 particles. 2D classification into 100 classes using default parameters were used to remove junk particles. The 214,619 particles contributing to the 2D classes bearing high-resolution features were used to compute two *ab-initio* reconstructions with 0 class similarity, initial mini batch size:300, final mini batch size 1000, initial resolution set to 20 Å and final resolution of 8 Å . Heterogeneous refinement into four classes with force hard classification “true” and 2000 batch size per class was performed on all 605,114 particles. The class with the best-resolved structural features, comprising 197,711 particles, was subjected to non-uniform refinement with C1 symmetry, an initial low-pass resolution set to 8 Å, minimize over per particle scale set to true, resulting in an aldolase reconstruction with a resolution of 6 Å . Particles were extracted with full box size (195,442 particles) and subjected to non-uniform refinement, followed by subsequent global and local CTF refinement followed by another round non-uniform refinement with C1 symmetry, yielding a reported global resolution of 3.3 Å based on the FSC 0.143 criterion (**Figs. S5, S6**). Duplicate particles were removed, resulting in 179,416 particles that were used for a non-uniform refinement with D2 symmetry imposed, an initial low-pass resolution set to 8 Å, and minimize over per particle scale set to “true.” The resulting structure had a reported global resolution of 3.1 Å based on the FSC 0.143 criterion.

#### Transthyretin

The image processing pipeline for TTR in the presence of DM is shown in **Fig. S7**. A total of 9,183 movies were collected and processed in cryoSPARC (v4.6-7). After motion correction and CTF estimation, micrographs with CTF fits worse than 6 Å and astigmatism above 600 Å were discarded. A total of 8,629 micrographs (post-processed) were used to extract around 6 million particles using 60-90 Å blob-picker in a 256-pixel box size, Fourier cropped to 100-pixel box size, which were subjected to 2D classification with 70 Å particle diameter and mask diameter of 75-90 Å, 40 online-EM iterations, and a batch size per class of 400. 1.5 M particles were selected and two *ab-initio* reconstructions were calculated, which were used in the heterogeneous refinement as initial volume, with a batch size per class of 2,000 and a spherical mask diameter of 100 Å . 858K particles were selected and further refined with non-uniform refinement with a 8 Å lowpass-filtered input and minimize over per-particle scales enabled, and re-extracted using the full 256-pixel box-size with recentering enabled. A second round of heterogeneous refinement was used to remove additional low-quality particles, resulting in 415K particles, which were refined in another non-uniform refinement. 3D classification ran in input mode, using a 4 Å filtered resolution with four classes. 388K particles corresponding to the highest-resolution 3D class were subjected to a local refinement with recentering using a mask covering the entire complex, resulting in a asymmetric and D2-symmetric final 3D reconstructions with a reported global resolutions of 3.3 Å and 3.2 Å, respectively (FSC cutoff at 0.143).

### Cryo-electron tomography of HA grids

Details are summarized in **Table S1**. Grids prepared with and without the detergent were used for cryoET analysis. For the sample with detergent addition immediately prior to grid freezing, concentrated sample of HA 10 mg/ml was mixed with DM dissolved in MQ at a final concentration of 0.06%, 3.5 uL of sample was deposited on glow discharged 300 mesh R1.2/1.3 UltrAuFoil Holey Gold Films (Quantifoil), blotted for 5 s after liquid spot on filter paper stop spreading and plunged frozen manually into liquid ethane. For grids without detergent 3mg/ml sample was used and grids were frozen in similar way as for DM sample. The collection was done using PACEtomo1.5^44^ in SerialEM ^45^ and 5 tomograms were collected on each grid. The magnification was 53,000X (pixel size of 1.6626 Å ) and a nominal defocus range between -4 and -6 µm with a 1-µm step. Tilt series collection was done in steps of 3° between -45° to 45° (total range of 90°) and a dose-symmetric scheme with 2.84 e/Å ^2^ per tilt (85.2 e/Å ^2^ total dose). Each tilt image was fractionated into 10 frames with the dose of 0.28 e/Å ^2^. The pre-processing and CTF estimation were done in WarpTools ^46^ and the tilt-series alignment was done using patch-tracking in Etomo ^47,48^. The alignment was then imported back into WarpTools and tomograms and their half-maps were reconstructed and Noise2Map was used for denoising ^49^.

### Atomic model building and refinement

For the NPM1, the cryo-EM structure PDB:8AS5 was used as the starting model ^21^. For the asymmetric Aldolase PDB:6V20 served as a starting model ^50^. The models were docked into the 3D reconstruction using UCSF ChimeraX ^42^. Manual model building and real-space structural refinement were performed with Coot ^51^ and Phenix ^52^, respectively. Phenix real-space refinement included global minimization, rigid body, local grid search, atomic displacement parameters, and morphing for the first cycle. It was run for 100 iterations, five macro-cycles, with a target bond RMSD of 0.01 and a target angle RMSD of 1.0. The refinement settings also include secondary structure restraints and Ramachandran restraints. ChimeraX plug-in ISOLDE ^53^ was used to manually fix any Ramachandran outliers, rotamers, and clashes not fixed by Phenix. Figures for publication were prepared using UCSF ChimeraX. Supplementary **Table S3** includes detailed parameters for the data collection, refinement, and validation statistics.

## Supporting information

Movie S1: HA without DM

Movie S2: HA with DM

## Authorship contributions

Maria Rafiq: Conceptualization, methodology, performed all cryo-EM structure determination, model building, refinement and deposition, Writing - original draft, review & editing. Jan-Hannes Schäfer performed TTR experiments, writing - review & editing. Hamidreza Rahmani: performed cryo-ET experiment, writing - review & editing, Shaochen You: expression and purification of NPM1, Michael Bollong: Writing - review & editing, Danielle Grotjahn: Writing - review & editing, Luke Wiseman: Writing - review & editing, Gabriel C. Lander: Writing - review & editing, Supervision, Project administration, Funding acquisition.

## Acknowledgments

We thank J.C. Ducom from Scripps High Performance Computing for computational support, W. Lessin at the Scripps Electron Microscopy facility for microscope support. We thank Julianna Han and Alesandra Rodriguez for providing HA (Strain A/California/04/2009 (H1N1, E47K HA2 stabilizing mutation), Jeff W. Kelly for providing TTR. This work was funded by the National Institute of Health grant GM143805. J.H.S. is supported by the German Research Council project number 556478029. M.R. is supported by the George E. Hewitt Foundation for Medical Research. D.A.G. is supported by the National Institutes of Health (NIH) grant GM154216-02.

## Data availability

The atomic coordinates for the NPM1, Aldolase (C1) and Aldolase (D2) structure have been deposited in the Protein Data Bank (PDB) under accession codes PDB:11IJ, PDB:10DS and 10IM respectively. The corresponding EM density maps (final unsharpened and sharpened maps, half maps, and masks) have been deposited to the Electron Microscopy Data Bank under accessions EMD-75715 for NPM1, EMD-75098 for asymmetric Aldolase, EMD-75198 for D2 symmetric Aldolase, EMD-75047 for asymmetric HA, and EMD-75400 for C3 symmetric HA respectively. The PDB and EMDB for TTR (EMD-74011 and 9ZBV), hClpP tetradecamer bound to bortezomib: (EMD-47234 and 9DW3).

## Competing interests

The authors declare that they have no known competing financial interests or personal relationships that could have appeared to influence the work reported in this paper.

## Supporting Information

**Figure S1.**
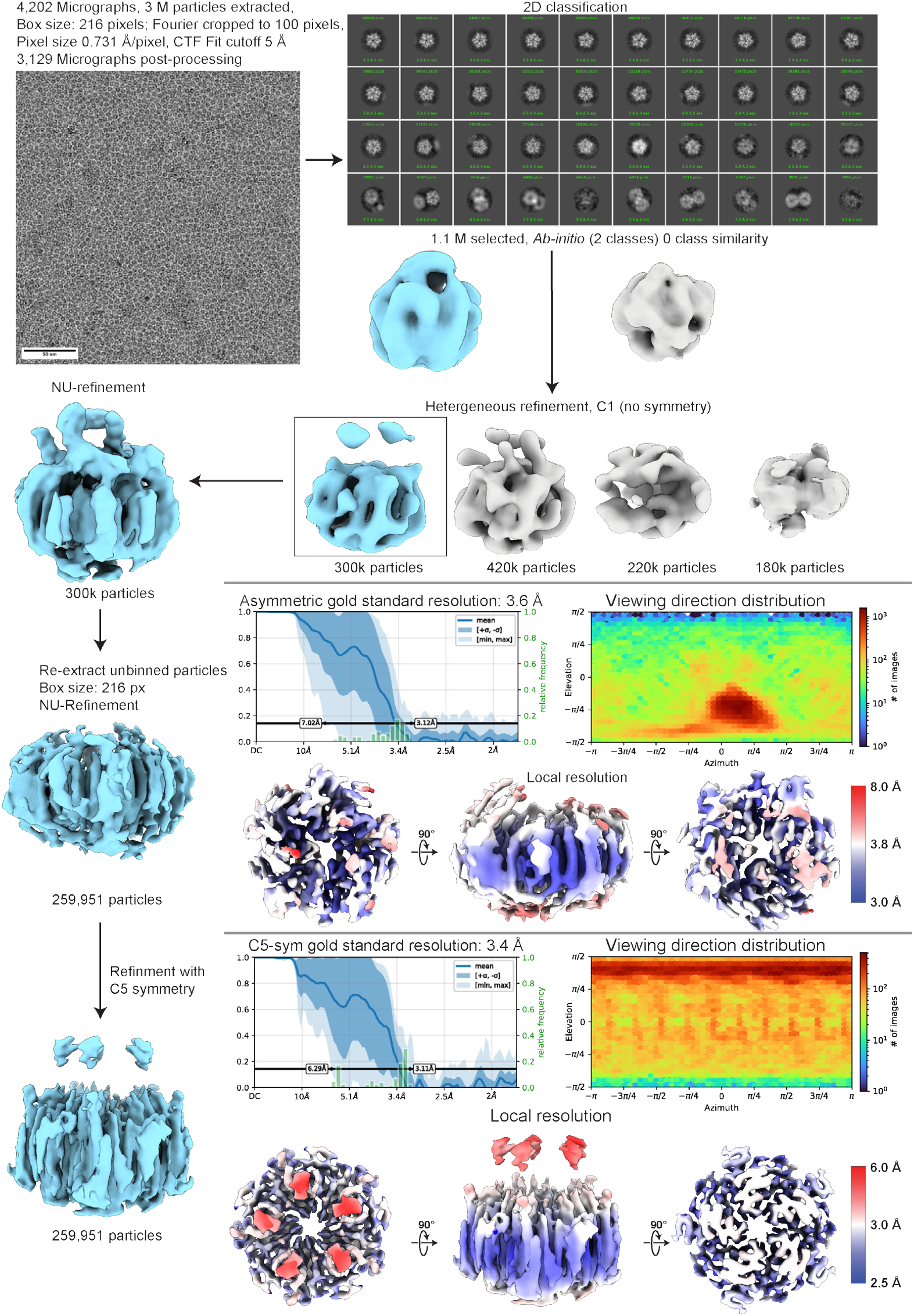
Cryo-EM image-processing workflows for NPM1 in the absence of detergent. A representative micrograph and 2D class averages are shown. Selected particles pertaining to structurally detailed classes were used to generate *ab-initio* reconstructions (two classes) and subsequently refined by heterogeneous refinement into four classes to isolate a well-resolved class. Particles from the well-resolved heterogeneous-refinement class were then re-extracted to full box size and refined with non-uniform refinement without symmetry and subsequently with imposed C5 symmetry. The cFAR and viewing direction distribution is shown for both reconstructions, along with the local resolution map below. Notably, the reconstruction exhibits severely anisotropic directional resolution.

**Figure S2.**
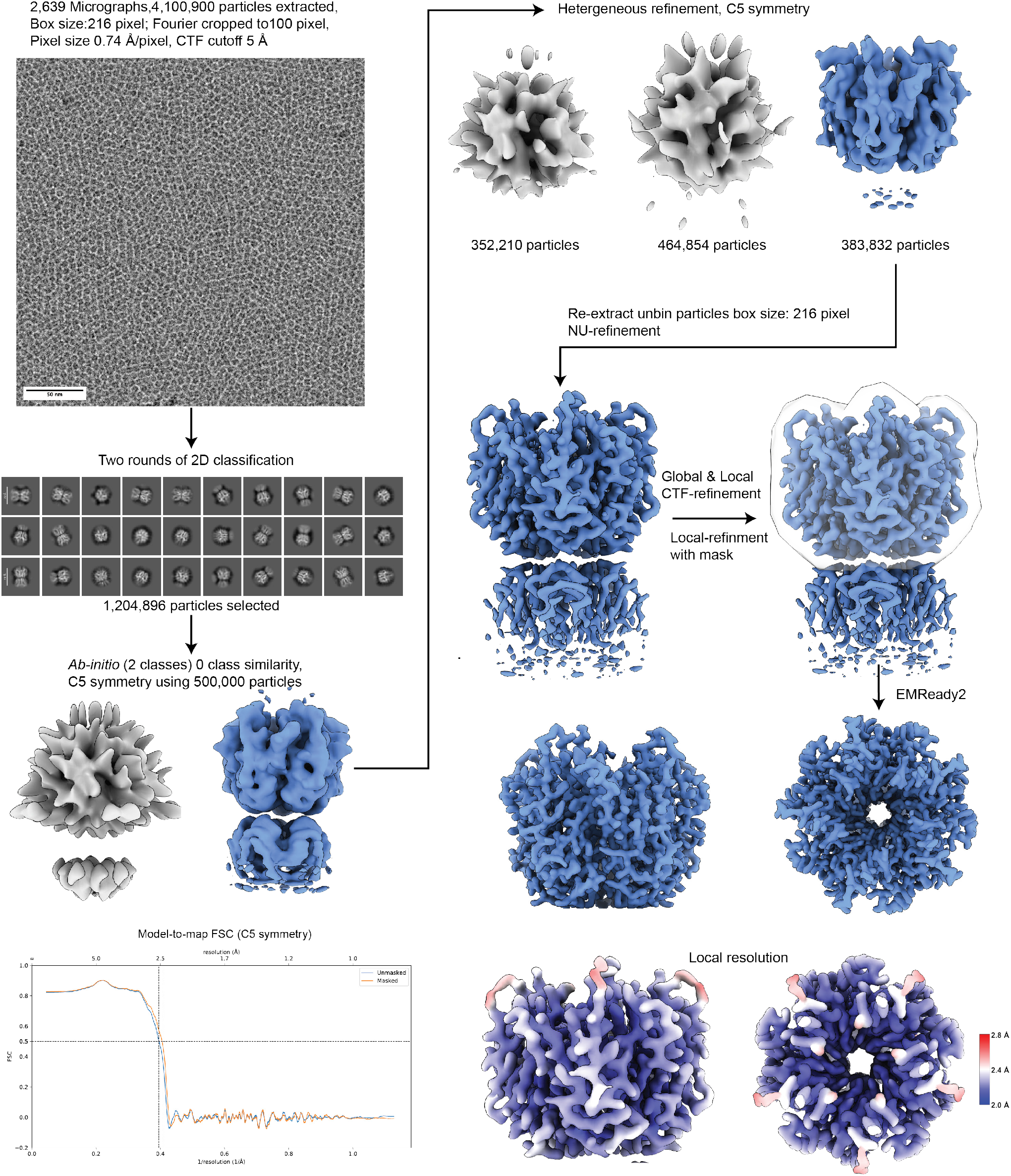
Cryo-EM image-processing workflows for NPM1 in the presence of DM. A representative micrograph and 2D class averages are shown. Selected particles pertaining to structurally detailed classes were used to generate *ab-initio* reconstructions (two classes) and subsequently refined by heterogeneous refinement into three classes to isolate a well-resolved class. Particles from the well-resolved heterogeneous-refinement class were then re-extracted to full box size and refined by non-uniform refinement, followed by global and local CTF refinement and masked local refinement. The mask is shown as a transparent volume. The final reconstruction with C5 symmetry imposed was reported with a global resolution of 2.4 Å, and was post-processed with EMReady2^54^The model-to-map FSC, with the cutoff at 0.5 denoted, and the local resolution map (estimated in Å ), are shown.

**Figure S3.**
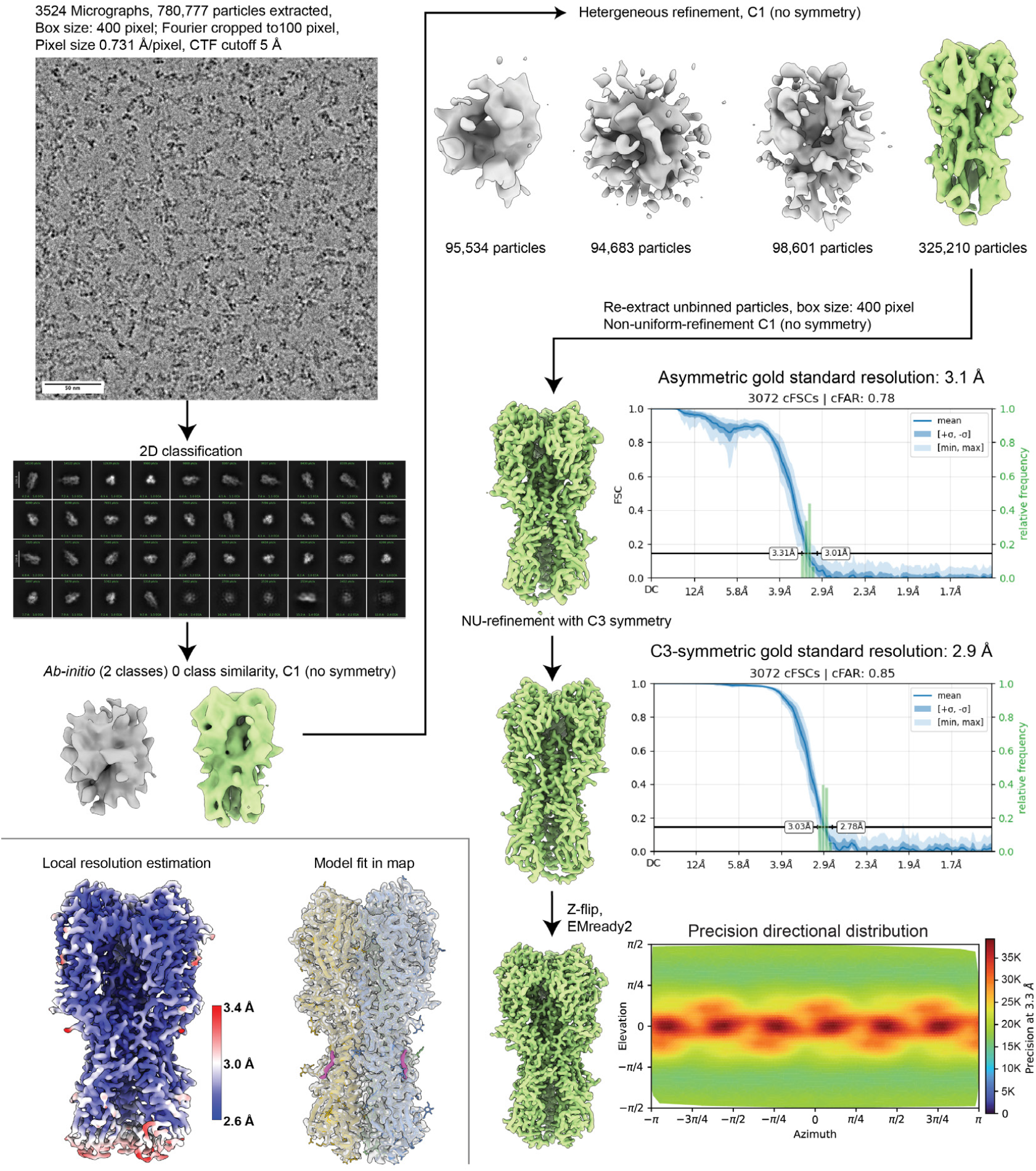
Cryo-EM image-processing workflows for HA in the presence of DM. A representative micrograph and 2D class averages are shown. Selected particles pertaining to structurally detailed classes were used to generate *ab-initio* reconstructions (two classes) and subsequently refined by heterogeneous refinement to four classes. Particles from the well-resolved heterogeneous-refinement class were then re-extracted to the full box size and refined by non-uniform refinement without symmetry and subsequently with imposed (C3) symmetry, yielding the final reconstructions reported to 3.1 Å and 2.9 Å resolutions, respectively. The cFAR plot is shown for both reconstructions, and the viewing direction distribution is shown for the C3-symmetric map. The final C3 map was post-processed with EMReady2. The map colored by the local resolution estimation is shown alongside a transparent map with an atomic model fit to show the quality throughout the structure. The densities corresponding to DM are colored magenta.

**Figure S4.**
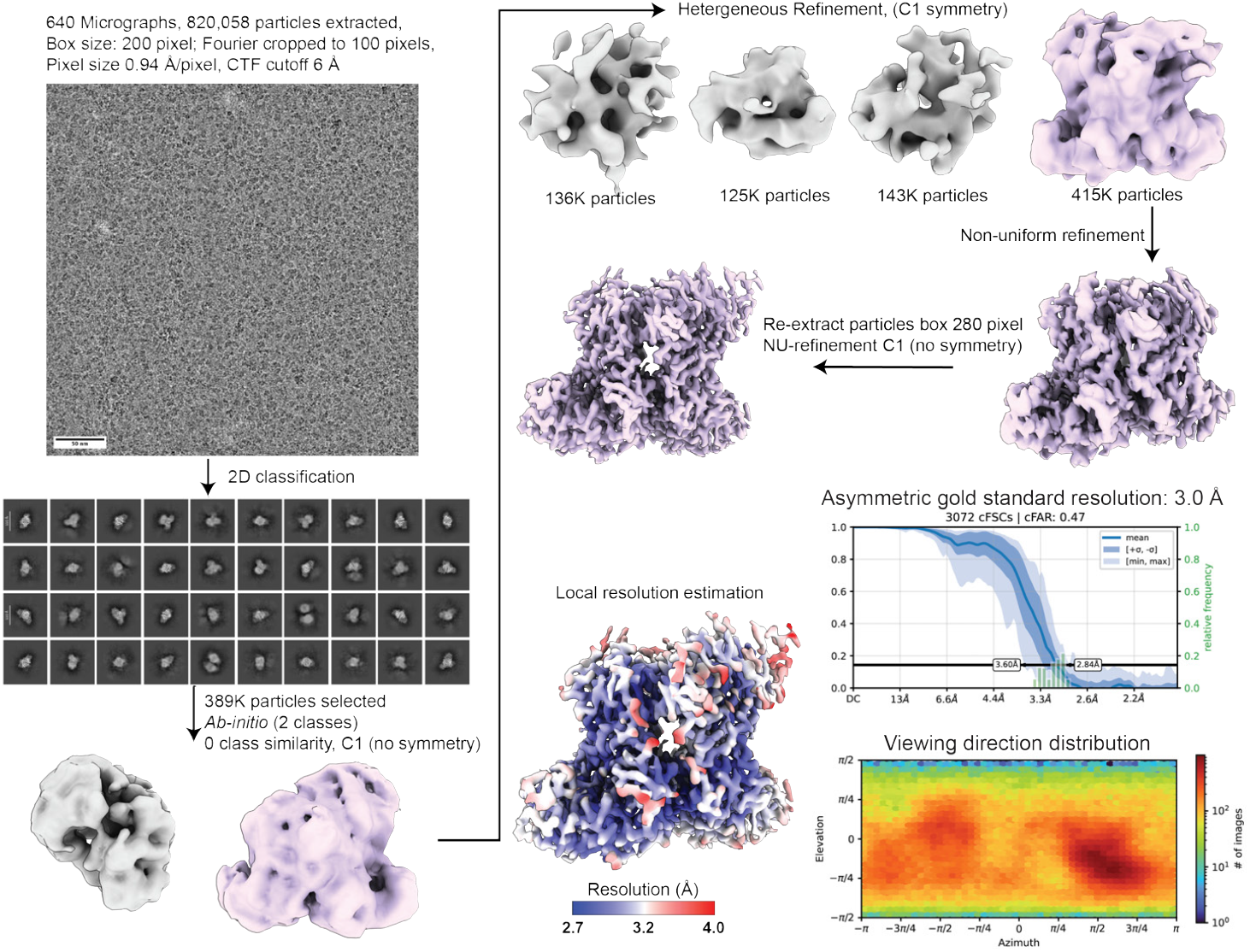
Cryo-EM image-processing workflows for asymmetric aldolase in the absence of detergent. A representative micrograph and 2D class averages are shown. Selected particles pertaining to the structurally detailed classes were used to generate *ab-initio* reconstructions (two classes) and subsequently refined with heterogeneous refinement to four classes. Particles from the well-resolved heterogeneous-refinement class were then re-extracted to full box size and refined by non-uniform refinement without symmetry, yielding the final reconstruction with a reported resolution of 3.0 Å . The cFAR and viewing direction distribution of the asymmetric reconstruction is shown below, alongside the map colored by local resolution.

**Figure S5.**
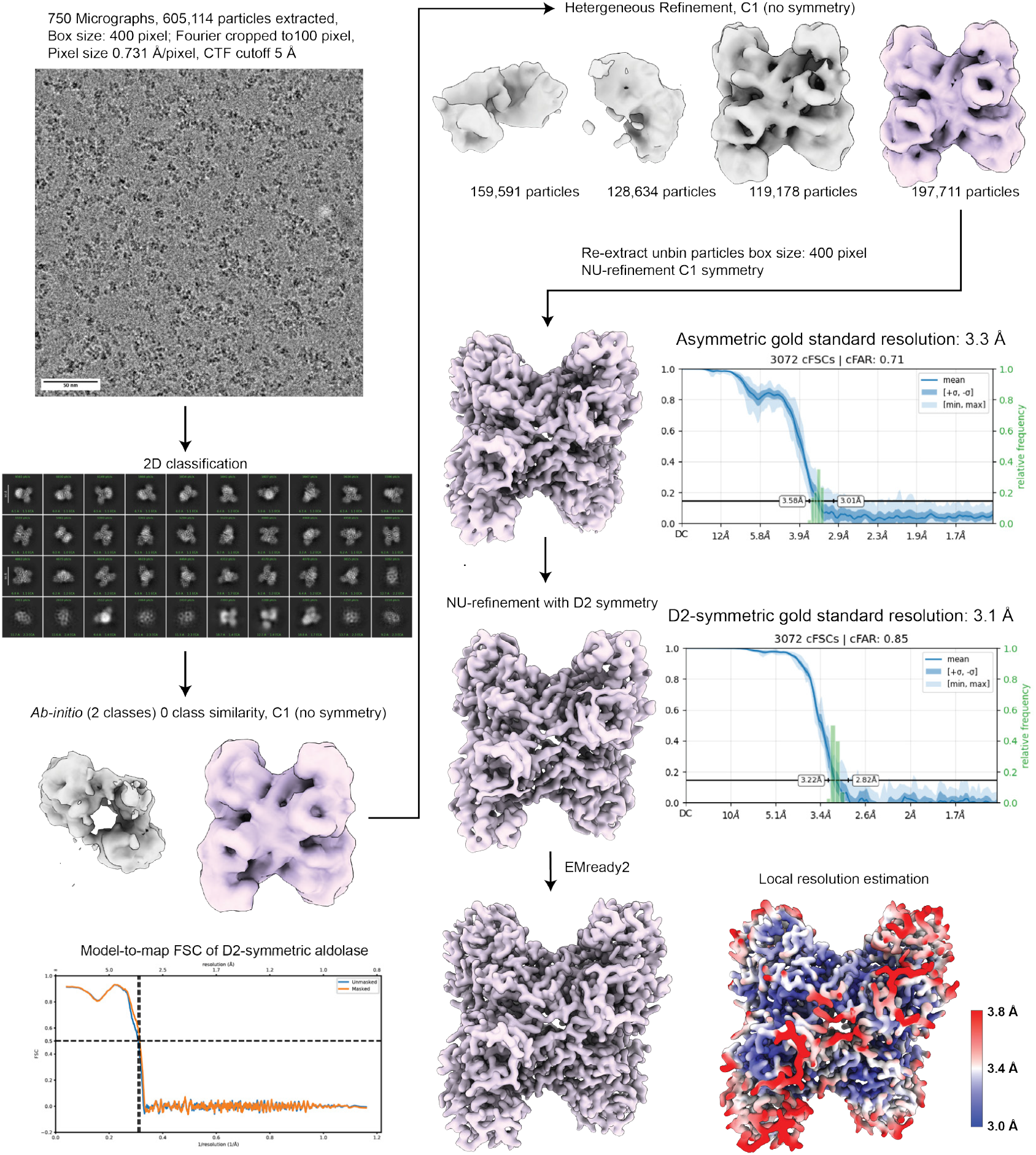
Cryo-EM image-processing workflows for aldolase in the presence of DM. A representative micrograph and 2D class averages are shown. Selected particles pertaining to the structurally detailed classes were used to generate *ab-initio* reconstructions (two classes) and subsequently refined by heterogeneous refinement to four. Particles from the well-resolved heterogeneous-refinement class were then re-extracted with a full box size and refined by non-uniform refinement without symmetry imposed and subsequently with D2 symmetry, resulting in maps with reported resolutions of 3.3 Å and 3.1 Å, respectively. The final map was post-processed with EMReady2^54^. The model-to-map FSC, with the cutoff at 0.5 denoted, is shown on the left, and the local resolution map (estimated in Å ), is on the right.

**Figure S6.**
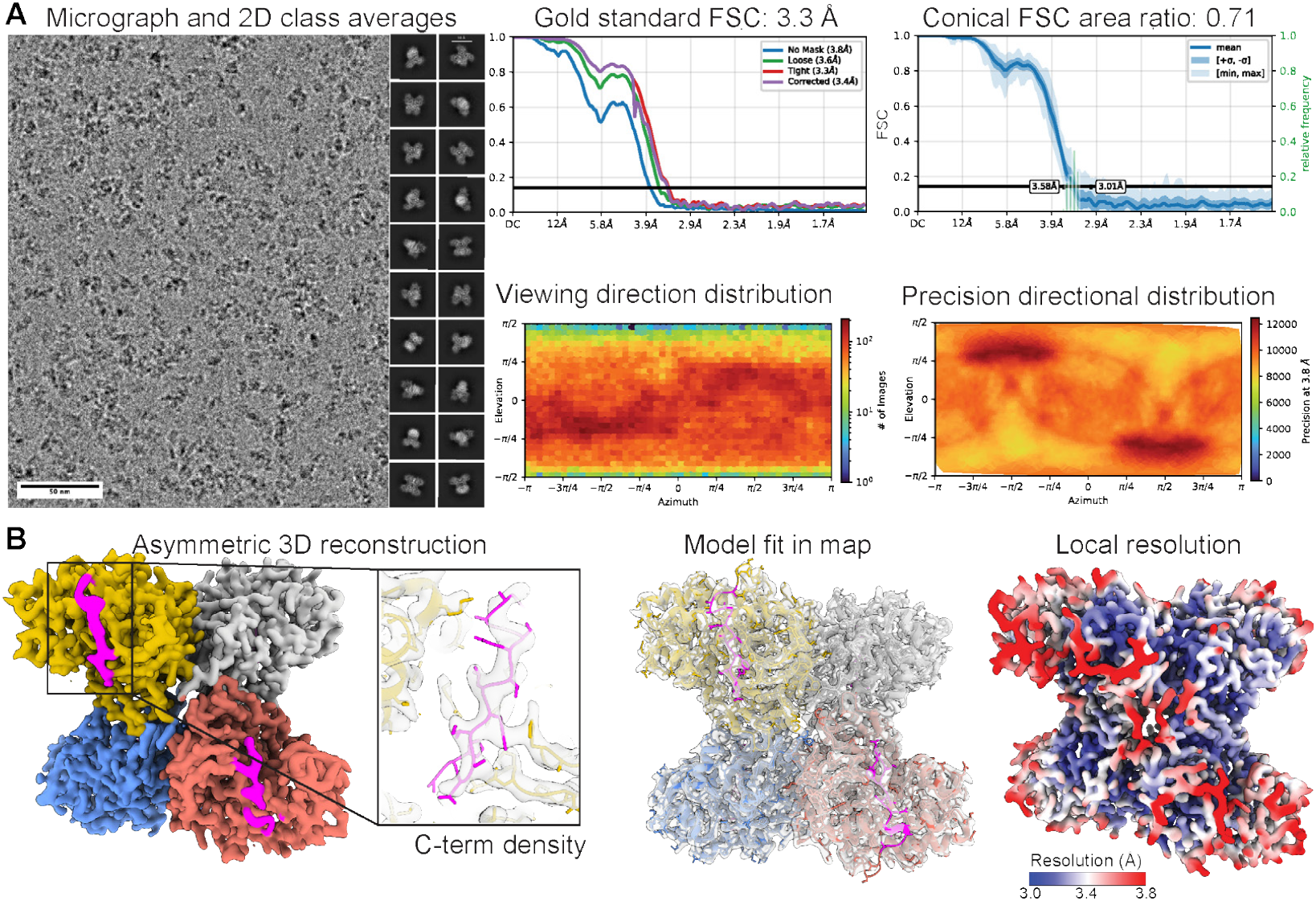
Asymmetric reconstruction of aldolase in the presence of DM. A representative micrograph and 2D class averages is shown. The gold-standard FSC plot reports an overall resolution of 3.3 Å at a 0.143 cutoff. The conical FSC plot with a cFAR of 0.71 indicates nearly isotropic directional resolution. The viewing-direction distribution and precision-directional distributions (below) reveal broad angular sampling. The resulting 3D reconstruction (far left panel) is shown (surface colored by each protomer), with density corresponding to the C-termini colored magenta. A close-up of the C-terminus is shown on the right, with residues fit into transparent EM density. The asymmetric map density (gray map) and fitted atomic model (colored by each protomer) is shown alongside the map colored by local resolution (estimated in Å ).

**Figure S7.**
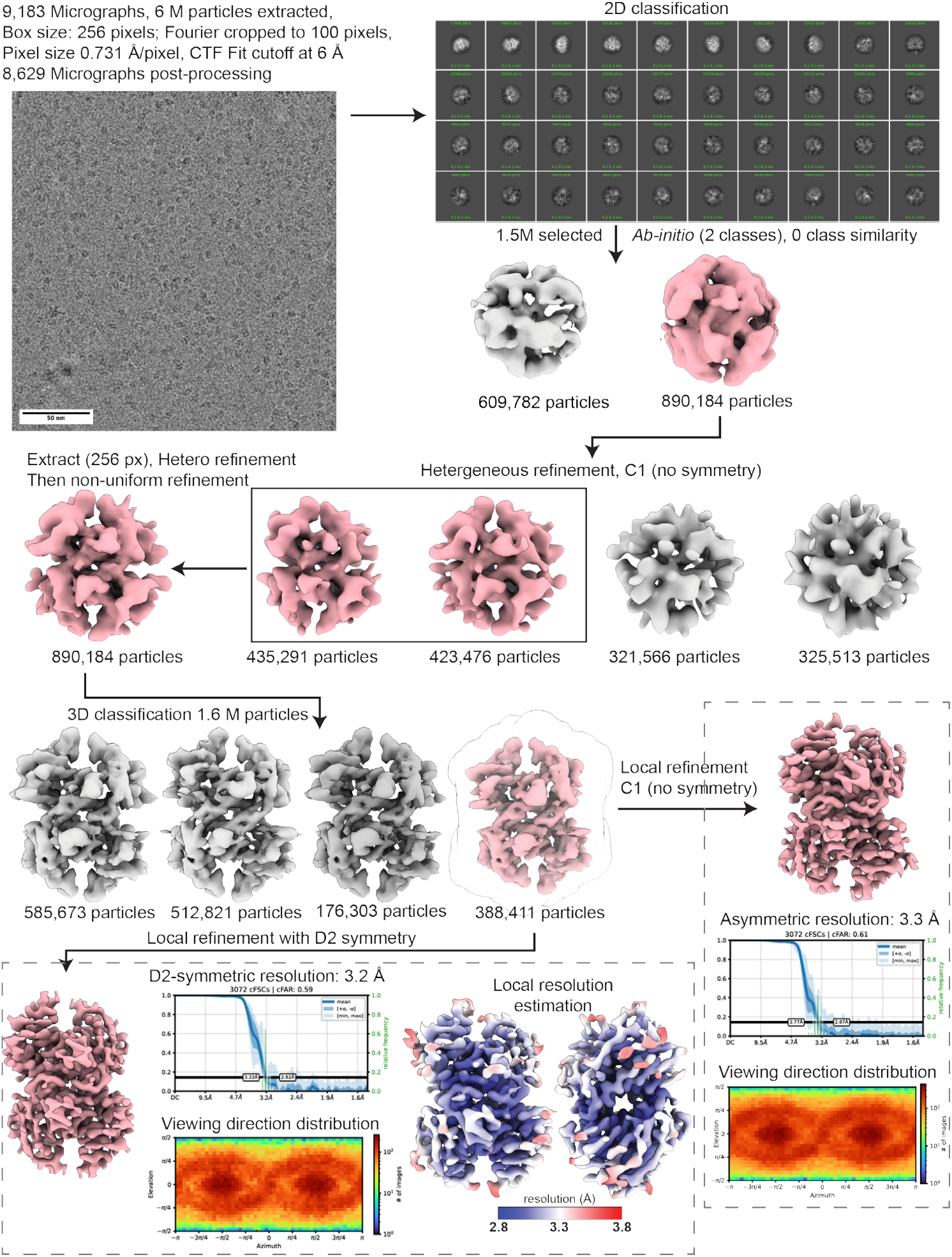
Cryo-EM image-processing workflows for TTR in the presence of DM. A representative micrograph and 2D class averages are shown. Selected particles from good classes were used to generate *ab-initio* reconstructions (two classes) and subsequently refined by heterogeneous refinement in four classes to isolate a well-resolved class which was further refined by non-uniform refinement. Particles were then re-extracted to full box size and refined by another round of hetero and non-uniform refinement with C1 symmetry. D2 symmetry was imposed to expand particles, further classified by 3D classification and class with best resolution was locally refined with and without symmetry, yielding the final reconstruction 3.2 Å and 3.3 Å respectively. Global resolution, cFAR and viewing direction distribution is shown for both asymmetric and symmetric reconstructions. Local-resolution estimation in Å is shown.

**Figure S8.**
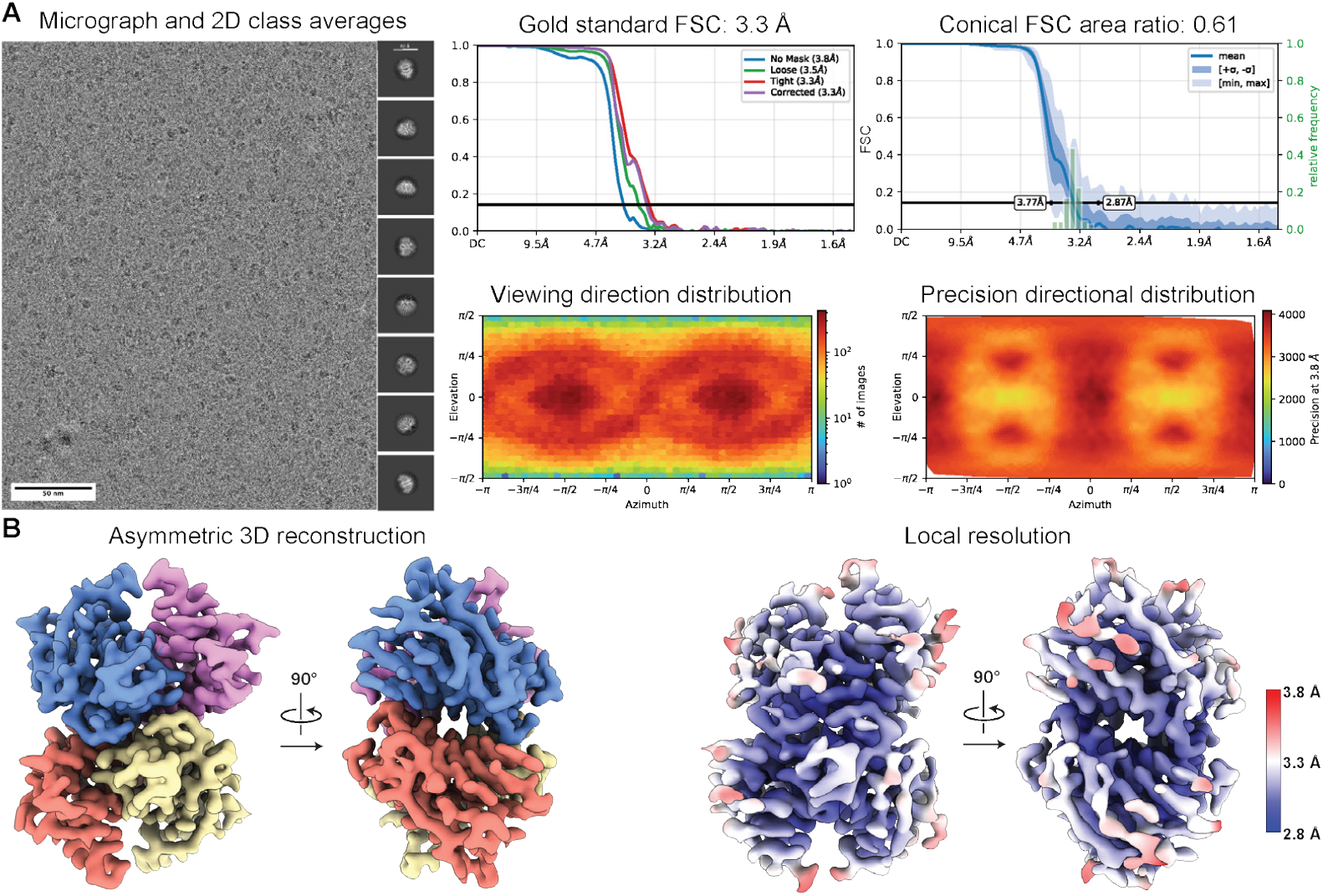
Asymmetric reconstruction of TTR in the presence of DM. A representative micrograph and 2D class averages are shown. The gold-standard FSC curve reports an overall global resolution of 3.3 Å at the 0.143 cutoff. The conical FSC (right panel) shows a cFAR of 0.61. The viewing-direction distribution and precision-directional distributions (center panels) reveal angular sampling with moderate preferential orientation, consistent with the cFAR value. The resulting 3D reconstruction (bottom left panels) is shown (surface colored by each protomer) alongside the map colored by local resolution (bottom right).

**Table S1.** NPM1 screening conditions used in this study.

| **Protein conc. (mg/mL)** | **Grid type** | **Vitrification** | **Detergent** | **Conc. (mM)** | **Type** |
| --- | --- | --- | --- | --- | --- |
| 2.5 / 5.0 | Open hole | Manual | LMNG | 0.01 | Non-ionic |
| 2.5 / 5.0 | Open hole | Manual | DDM | 0.17 | Non-ionic |
| 2.5 / 5.0 | Ultra-thin carbon | Manual | LMNG | 0.01 | Non-ionic |
| 10 | Open hole | Manual | FC | 3 | Zwitterionic |
| 10 | Graphene | Manual | GDN | 0.018 | Non-ionic |
| 10 | Graphene | Manual | CHAPSO | 4 | Zwitterionic |
| 8 | Open hole | Manual | CHAPSO | 4, 2 | Zwitterionic |
| 20 | Graphene + Poly-lysine | Manual | FC | 0.75 | Zwitterionic |
| 0.25 | Graphene | Manual | None | None |  |
| 0.25 | Graphene affinity grids | Vitrobot | None | None |  |
| 10 | Open hole | Manual | DM | 1.8 | Non-ionic |

**Table S2.** SPA Cryo-EM data collection parameters and image processing of NPM1, Aldolase, and HA. (NA: not applicable)

|  | **NPM1** | **Aldolase (C1)** | **Aldolase (D2)** | **HA (C1)** | **HA (C3)** |
| --- | --- | --- | --- | --- | --- |
| **EMDB ID** | EMD-75715 | EMD-75098 | EMD-75198 | EMD-75047 | EMD-75400 |
| **PDB ID** | 11IJ | 10DS | 10IM | NA | NA |
| **Data Collection** |  |  |  |  |  |
| Microscope | Arctica | Arctica | Arctica | Arctica | Arctica |
| Camera | Falcon 4i | Falcon 4i | Falcon 4i | Falcon 4i | Falcon 4i |
| Magnification (nominal) | 190,000 | 190,000 | 190,000 | 190,000 | 190,000 |
| Voltage (keV) | 200 | 200 | 200 | 200 | 200 |
| Total dose (e<sup>-</sup>/Å<sup>2</sup>) | 50 | 50 | 50 | 50 | 50 |
| EER Fractions | 40 | 40 | 40 | 40 | 40 |
| Pixel size (Å/pixel) | 0.74 | 0.731 | 0.731 | 0.731 | 0.731 |
| Defocus range (μm) | -2.0 to -0.8 | -2.0 to -0.8 | -2.0 to -0.8 | -2.0 to -0.8 | -2.0 to -0.8 |
| Recorded movies | 2639 | 750 | 750 | 3524 | 3524 |
| **Image Processing** |  |  |  |  |  |
| Processing software | CS (v4.6) | CS (v4.7) | CS (v4.7) | CS (v4.6) | CS (v4.6) |
| Final particle images | 383832 | 195442 | 179416 | 325210 | 325210 |
| Symmetry imposed | C5 | C1 | D2 | C1 | C3 |
| Resolution (FSC 0.143) (Å) | 2.41 | 3.36 | 3.10 | 3.10 | 2.95 |
| Local resolution (Å) | 2.0–2.8 | 3.0–3.8 | 3.0–3.8 | 2.8–3.8 | 2.6–3.4 |
| cFAR | 0.82 | 0.73 | 0.85 | 0.78 | 0.78 |
| Sharpening B-factor (Å<sup>2</sup>) | -93 | -109 | -153 | -115 | -129 |
| **Model Refinement** |  |  |  |  |  |
| Initial model | 8AS5 | 6V20 | 6V20 |  |  |
| FSC map-to-model (0.5) (Å) | 2.5 | 3.5 | 3.2 |  |  |
| MolProbity score | 0.84 | 1.48 | 1.36 |  |  |
| Clash score | 1.24 | 5.64 | 6.44 |  |  |
| **Composition** |  |  |  |  |  |
| Chains | 5 | 4 | 4 |  |  |
| Atoms | 7765 | 21872 | 21884 |  |  |
| Protein residues | 530 | 1436 | 1448 |  |  |
| **Bonds (RMSD)** |  |  |  |  |  |
| Length (Å) | 0.002 | 0.002 | 0.002 |  |  |
| Angles (°) | 0.441 | 0.442 | 0.386 |  |  |
| **B-factors (min/max/mean)** |  |  |  |  |  |
| Protein residues | 3.25/77.09/15.05 | 20.66/147.39/67.60 | 34.11/155.15/83.48 |  |  |
| **Ramachandran plot (%)** |  |  |  |  |  |
| Favored | 99.04 | 96.71 | 98.89 |  |  |
| Allowed | 0.96 | 3.29 | 1.11 |  |  |
| Outliers | 0.00 | 0.00 | 0.00 |  |  |
| Rotamer outliers (%) | 0.00 | 0.70 | 0.69 |  |  |
| Q-score | 0.81 | 0.67 | 0.71 |  |  |
| Ligand | NA | NA | NA |  |  |

**Table S3.** Microscope parameters for collection of cryo-ET data.

| Parameter | Value |
| --- | --- |
| Microscope | Titan Krios |
| Magnification | 53,000 |
| Camera | K3 |
| Voltage (kV) | 300 |
| Electron dose per image ($e^-/\text{\AA}^2$) | 2.84 |
| Fractions per image | 10 |
| Tilt range (increment) | $-45^\circ$ to $+45^\circ$ ($3.0^\circ$) |
| Defocus range ($\mu\text{m}$) | -4 to -6 |
| Pixel size ($\text{\AA}$) | 1.6626 |

## Appendix A.

**Supplementary data**

**Movie S1: HA trimer with no detergent**

**Movie S2: HA trimer with detergent**

## Notes

### Competing Interest Statement

The authors have declared no competing interest.

